# Chromatin 3D structure reconstruction with consideration of adjacency relationship among genomic loci

**DOI:** 10.1101/741447

**Authors:** Fang-Zhen Li, Zhi-E Liu, Xiu-Yuan Li, Li-Mei Bu, Hong-Xia Bu, Hui Liu, Cai-Ming Zhang

## Abstract

Chromatin 3D conformation plays important roles in regulating gene or protein functions. High-throughout chromosome conformation capture (3C)-based technologies, such as Hi-C, have been exploited to acquire the contact frequencies among genomic loci at genome-scale. Various computational tools have been proposed to recover the underlying chromatin 3D structures from in situ Hi-C contact map data. As connected residuals in a polymer, neighboring genomic loci have intrinsic mutual dependencies in building a 3D conformation. However, current methods seldom take this feature into account. We present a method called ShNeigh, which combines the classical MDS technique with local dependence of neighboring loci modelled by a Gaussian formula, to infer the best 3D structure from noisy and incomplete contact frequency matrices. The results obtained on simulations and real Hi-C data showed, while keeping the high-speed nature of classical MDS, ShNeigh is more accurate and robust than existing methods, especially for sparse contact maps. A Matlab implementation of the proposed method is available at https://github.com/fangzhen-li/ShNeigh.

**Author summary:** We propose a new method to infer a consensus 3D genome structure from a Hi-C contact map. The novelty of our method is that it takes into accounts the adjacency of genomic loci along chromosomes. Specifically, the proposed method penalizes the optimization problem of the classical multidimensional scaling method with a smoothness constraint weighted by a function of the genomic distance between the pairs of genomic loci. We demonstrate this optimization problem can still be solved efficiently by a classical multidimensional scaling method. We then show that the method can recover stable structures in high noise settings. We also show that it can reconstruct similar structures from data obtained using different restriction enzymes.

## Introduction

Correct 3D organization of chromosomes plays important roles in maintaining chromosomal functions such as gene expression, epigenetic modification and timely copy and separation of chromosomes in mitosis. However, determining chromosomal 3D structures is still an unsettled issue currently. Traditional techniques such as fluorescence microscope and fluorescence in situ hybridization (FISH), usually have low resolution and can only probe a few of individual genome loci at one time. Hi-C[1], which is derived from Chromatin conformation capture (3C) and depth sequencing technique, provides a new promise for this problem. As a high-resolution and high-throughout method of studying chromosomal 3D conformation, Hi-C can measure the contact frequency between genome loci pairs at the genome-wide level. Inferring the 3D structure of the genome from the contact frequency matrix obtained by Hi-C has become an interesting research topic of bioinformatics since the occurrence of Hi-C.

However, reconstructing the 3D structures of chromosomes from the Hi-C data is not so straightforward but an optimization problem essentially. As in other applications, a standard optimization procedure requires clarifying two issues: the objective function to be minimized or maximized and the optimization algorithm. As for the objective function, one strategy is the distance-based formula. That is, this strategy first converts the contact frequency matrix into the spatial distance matrix and then minimizes the discrepancy between the distance matrix calculated from the predicted structure and that converted from the frequency matrix [2,14-18]. Two operations are prerequisite for this strategy: first, the frequency matrix is normalized to remove the biases related to the DNA sequence, among which GC content, sequence mappability and frequency of restriction sites are three most apparent bias resources [4]; second, the conversion factor that modulates the power law relationship between the frequency matrix and the distance matrix [1] is estimated through an additional optimization procedure [2]. Another strategy of selecting the objective function casts the problem of structure inference as a maximum likelihood problem by assuming the contact frequency between genome loci follows a Poisson distribution [3,5]. HSA [11] constructs the likelihood by integrating multiple contact matrices generated from different enzymes. The advantage of this strategy is that, by modeling the effect of all the three data bias (i.e. GC content, sequence mappability and frequency of restriction sites) and the power law relationship between frequency and distance matrix with a generalized linear formula, all these effects can be absorbed into the final likelihood function. Thus, all parameters --- the Cartesian coordinates of all genome loci, the coefficients describing the effect of data bias and the conversion factor parameter --- can be derived simultaneously through a unified optimization procedure. Consequently, the normalization of the contact frequency matrix and the additional conversion factor inference procedure, which are requisite for the first strategy, are now unnecessary.

No matter which objective function above is adopted, the issue finally boils down to a nonconvex, nonlinear, large scale optimization problem, for which a simple local searching approach, such as Newton algorithm, is not suitable. Several global searching schemes have been proposed. ChromSDE[2] transforms the problem into a semi-definite programming (SDP) problem by embedding the original 3D Euclidean space into the Hilbert space of higher dimension. It can guarantee recovering the correct structure in the noise-free case. But for noisy input data a local optimization method is needed to refine the solution obtained from the SDP problem. PASTIS [3] uses IPOPT [7], a C++ package that implements an interior point filter algorithm for large-scale nonlinear optimization, to maximize the Poisson likelihood. BACH and BACH-MIX [5] apply Gibbs sampler with hybrid Monte Carlo to draw samples in the parameter space and output a collection of 3D chromosomal structures from the Bayesian posterior distribution. TADbit [8,9] contains a module of chromosome 3D reconstruction that was developed around Integrative Modeling Platform (IMP, http://www.integrativemodeling.org), a general framework for restraint-based modeling of 3D bio-molecular structures [10]. HSA [11] adopts simulated annealing combined with Hamiltonian dynamics to explore the chromatin comformal space. Different from BACH et al., MCMC5C [12] assumes the contact frequency is normally distributed and employs the Markov chain Monte Carlo (MCMC) with Metropolis-Hastings sampler [13] to sample from the posterior distribution. Same as BACH, MCMC4C outputs an ensemble of conformations. AutoChrom3D [14] selects LINGO (www.lindo.com/products/lingo), a commercial nonlinear constrained optimizer, to get the best chromatin structure. 3DMax [15] utilizes a stochastic gradient ascent algorithm to maximize the likelihood generated from the normal distribution. MOGEN [16-17] and LorDG [18] maximized the objective function by using steepest gradient ascent with the back-tracking line search algorithm.

The problem of inferring the coordinates of N objects in the 3D space from the distance information between them can be solved perfectly by the classical multidimensional scaling method (MDS) [6]. However, the distance matrix converted from the contact frequency matrix is not complete in that it contains many unknown entries generally, which makes the classical MDS method can not be utilized directly. This is just why various optimization approaches above mentioned were proposed. In order to avoid the time-consuming optimization procedure, ShRec3D [19] cleverly designed a two-step algorithm. It first completes the distance matrix by using the concept of shortest path in graph theory (i.e. Floyd-Warshall algorithm), and then exerts the classical MDS to reconstruct 3D genome structures. It is orders of magnitude faster than the above optimization-based methods. ShRec3D+ [20] corrects the conversion factor by a golden section search before carrying out ShRec3D. MDSGA [21] improves the shortest path distances using a genetic algorithm.

It should be noted that the positions of genomic loci in the 3D space are not irrelevant to each other. Genomic loci can be taken as a bunch of connected beads that comprise of a polymer. Two loci adjacent in the genome are surely close to each other in the 3D space. However, current methods seldom give consideration to this property of genomes. HSA [11] characterizes the adjacency relationship of neighboring loci by a Gaussian Markov chain to capture the local dependence of genomic loci. In the present work we extend the framework of classical MDS and provide a more flexible way to model the correlations between genomic loci of local proximity. Our algorithm, named ShNeigh, can significantly improve the performance of ShRec3D and simultaneously still runs far faster than the optimization-based methods, such as ChromSDE.

## Results

### Simulated data study

We compared our ShNeigh with the existing methods ChromSDE [2], ShRec3D [19] and ShRec3D+ [20]. As for ChromSDE, the quadratic SDP algorithm is adopted. We first test these programs on the simulated helix structure dataset. **Figure 3** shows the performance comparison for the programs under different measurements. We draw the mean result of 10 runs for each noise level to reduce the occasional fluctuation. The conversion factor is always assumed equal to 1 in ShRec3D, which is just the true value for our simulated data. Because ShRec3D+ merely adds a conversion factor estimation step upon ShRec3D and can not improve the performance of ShRec3D for the simulation scenario, it is not included in Figure 2a-b. As expected, when the noise level increases, SCC decreases and RMSD increases generally. The RMSD of ChromSDE starts from 0 at zero noise level, which coincides with the claim that ChromSDE can guarantee recovery of the true structure in the noise-free case. Unfortunately, the other three programs do not possess such a good feature. However, when the noise level get larger (>0.25), the superior behavior of ShNeigh1 and ShNeigh2 begins to emerge, and their superiority enlarges compared to ChromSDE with the increasing noise level (Figure 2a). ShNeigh1 and ShNeigh2 perform similarly and both significantly outperform ShRec3D, showing that inclusion of the neighboring dependency relationship can offer essential improvement against the underlying ShRec3D method. In summary, our ShNeigh algorithms are more robust and accurate than ChromSDE and ShRec3D, except for comparing to ChromSDE in the noise-free or little noise situation. However, Figure 2b shows ShNeigh1 and ShNeigh2 have no pronounced improvement against ShRec3D in terms of the SCC measure, and both ShRec3D and ShNeigh programs perform worse than ChromSDE on SCC. It seems that ChromSDE tends to be over faithful to the noisy input data, which may be the reason why ChromSDE is less robust than other programs.

**Fig 1.**
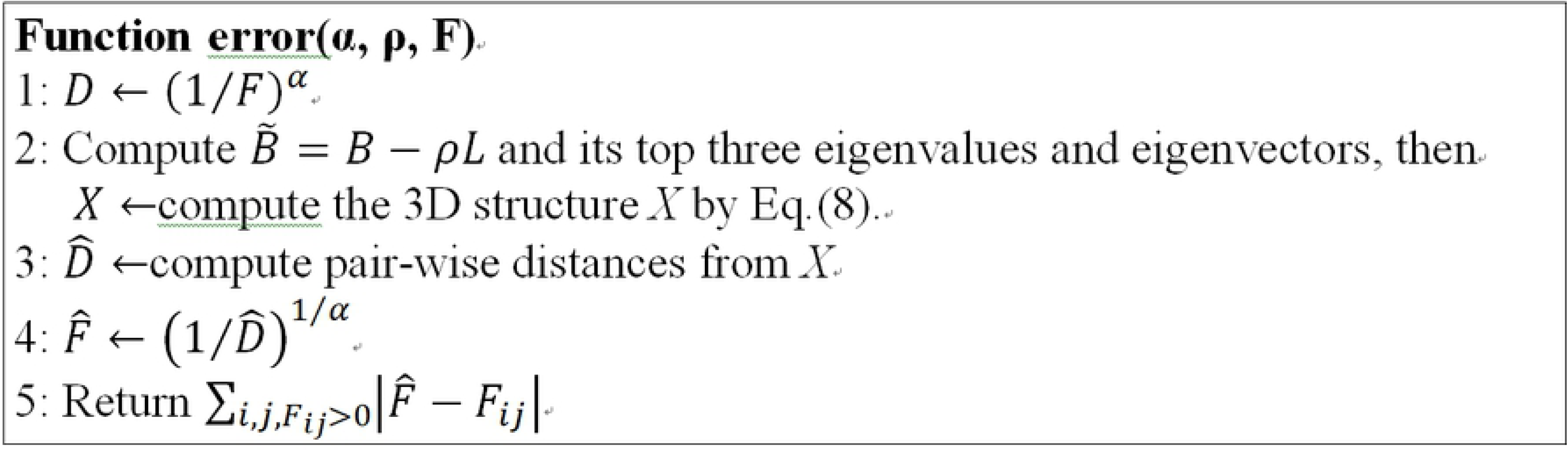
Goodness function definition.

**Fig 2.**
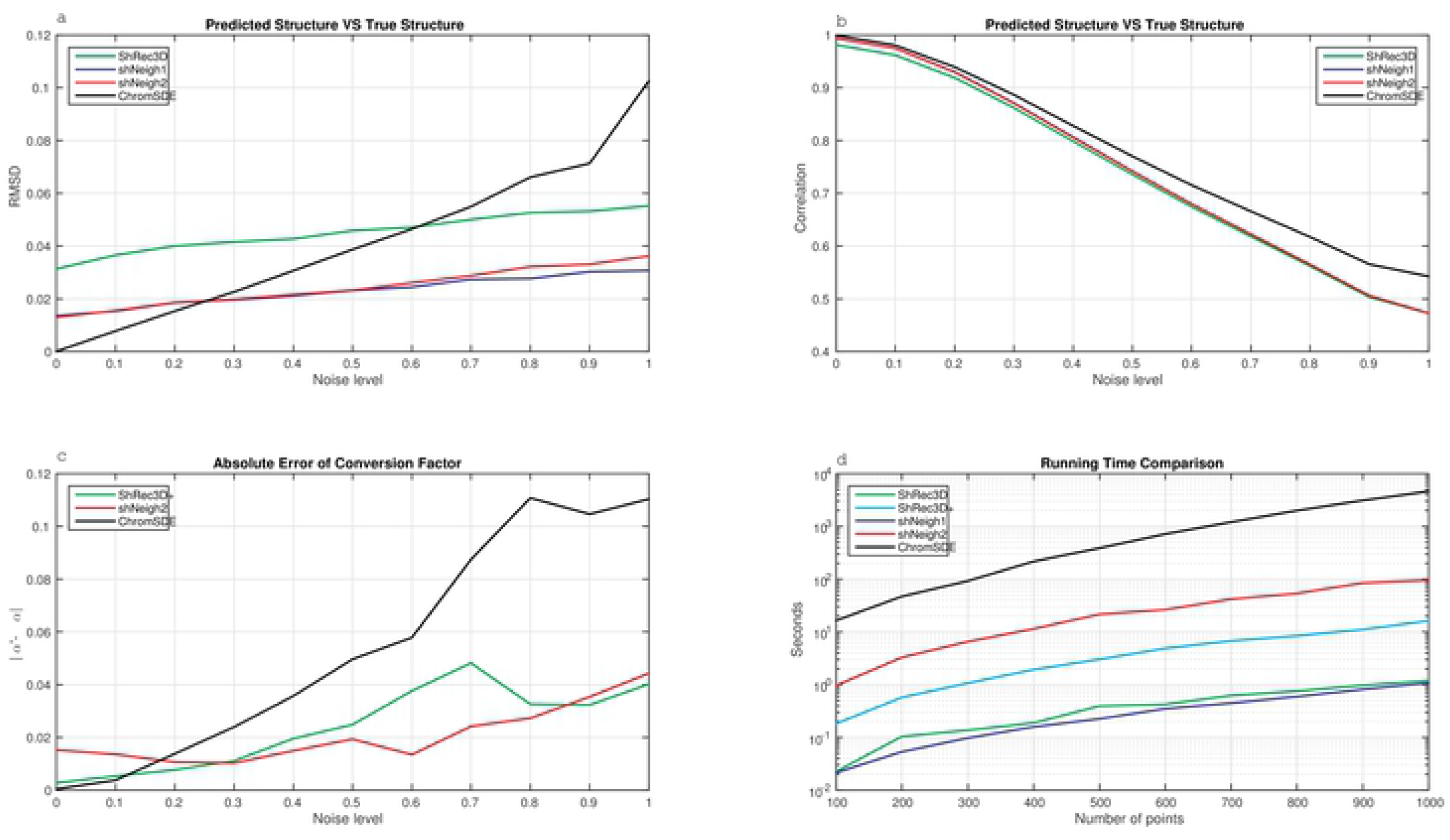
Performance comparison on simulated data. **(a)** Spearman correlation between the distance matrices calculated from the predicted structure and those from the true structure under varying noise levels. **(b)** Root mean square deviation (RMSD) between the predicted structure and the true structure under varying noise levels. **(c)** The absolute difference between the estimated and true conversion factor under varying noise levels. (d) Logarithm of running time of tested programs under varying number of points.

**Fig 3.**
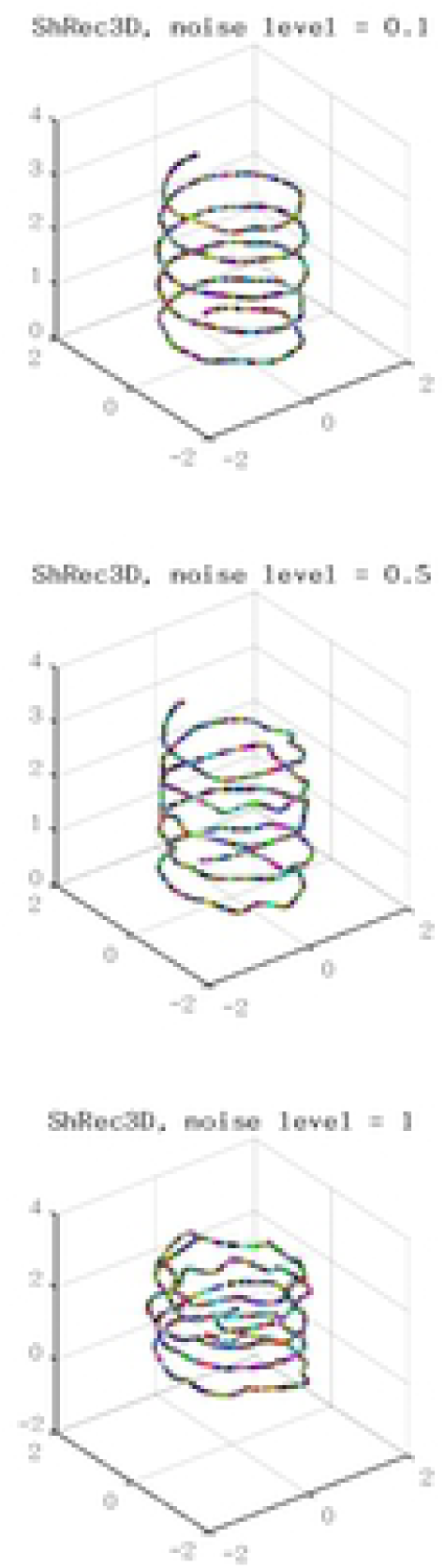
3D Structures predicted by different methods on simulated helix data under different noise levels. The red curve is the true structure and the green curve is the predicted structure. ShNeigh uses ShNeigh1, and ShNeigh2 has similar performance.

In Figure 2c the absolute error between the estimated conversion factor α and the true α (=1) rises with increasing noise level generally. At low noise levels (<0.2), ChromSDE can nearly perfectly estimate α values, consistent with its performance on the RMSD measure. But when the noise level increases the error estimated by ChromSDE ascends dramatically, indicating ChromSDE is prone to give a wrong conversion factor estimation as the data get more noisy. By contrast, ShNeigh2 and ShRec3D+ can estimate the conversion factor α quite accurately across various noise levels. We can see from Figure 2a,c that the performance of these programs on RMSD and that on the absolute α error interweave with each other, in that samller RMSD leads to smaller α error, and vice versa.

As described in the previous section, ShRec3D and ShNeigh1 have no optimization, while ShRec3D+ includes a uni-variate optimization step (estimate α) and ShNeigh2 possesses a two-variate minimum searching procedure (estimate α and the weight ρ). Therefore, ShRec3D and ShNeigh1 are most efficient among the tested programs, and ShRec3D+ runs slower than ShRec3D and ShNeigh1 and faster than ShNeigh2 (Figure 2d). ChromSDE is the most time-consuming since it needs to explore a space of *N*^2^ variables (compute a semi-definite kernel matrix).

Figure 3 shows the predicted structures of the simulated helix by different programs (ShRec3D, ShNeigh2 and ChromSDE) under different noise levels. The structures predicted by ShNeigh1 are very similar to ShNeigh2 and so not shown. For the noise-free case drawn in the top row, both ShNeigh2 and ChromSDE can almost perfectly recover the true structure, and ShRec3D seems to give a bit over-fat structure. For the case of medium noise level (=0.5, the middle row), the performances of all the three programs get worse, but the reconstruction result of ShNeigh2 is still quite good, and it is difficult to identify the helix structure from ChromSDE’s reconstruction. When the noise level gets the maximum (=1, the bottom row), ShNeigh2 can still present a clear helix structure, and by contrast, the structure by ShRec3D is too fat and obscure, while ChromSDE completely fails. We conclude that, on the whole, ShNeigh outperforms ShRec3D and ChromSDE, especially in the highly noisy circumstance.

At last we investigate the impact of signal coverage on the performance of these programs. Obviously signal coverage is proportional to the number of nearest neighbors parameter *K* of the simulation code. In fact signal coverage is approximately equal to *K*/*N* (Figure 4f). Figure 4 shows RMSD increases with descending nearest neighbors *K* for all programs and all noise levels, indicating that reducing signal coverage can substantially deteriorate the reconstruction results. Our programs ShNeigh1 and Sheigh2 perform similarly and both of them give apparent improvement relative to ShRec3D for all noise levels and all signal coverage. And they outperform ChromSDE at most situations. It is only at low noise level or high signal coverage that ChromSDE performs better than ShNeigh1 and ShNeigh2 (Figure 4a-b). The leading status of ShNeigh1 and ShNeigh2 compared to ChromSDE gets more significant when the noise level increases, which coincides with the results shown in Figure 2-3. When the signal coverage decreases, ChromSDE’s RMSD gets larger rapidly, while our ShNeigh programs are less sensitive to the signal sparseness. Therefore, the leading status of ShNeigh1 and ShNeigh2 compared to ChromSDE also gets more significant when the frequency matrix turns sparser (Supplementary Figure S1).

**Fig 4.**
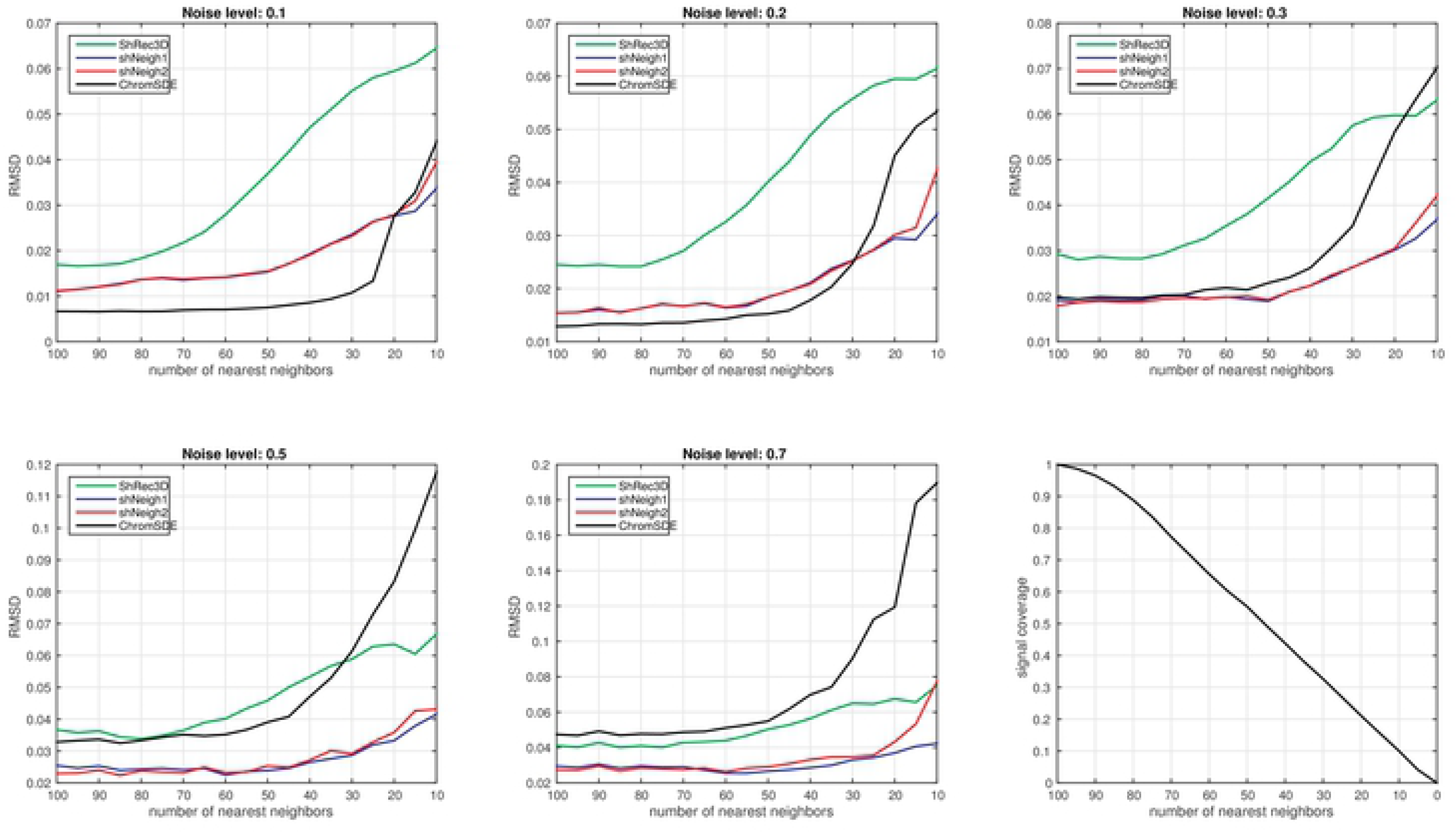
RMSD measure of tested programs on simulated data under varying number of nearest neighbors *K*. The point number of the simulated helix is 100.

**Fig.5.**
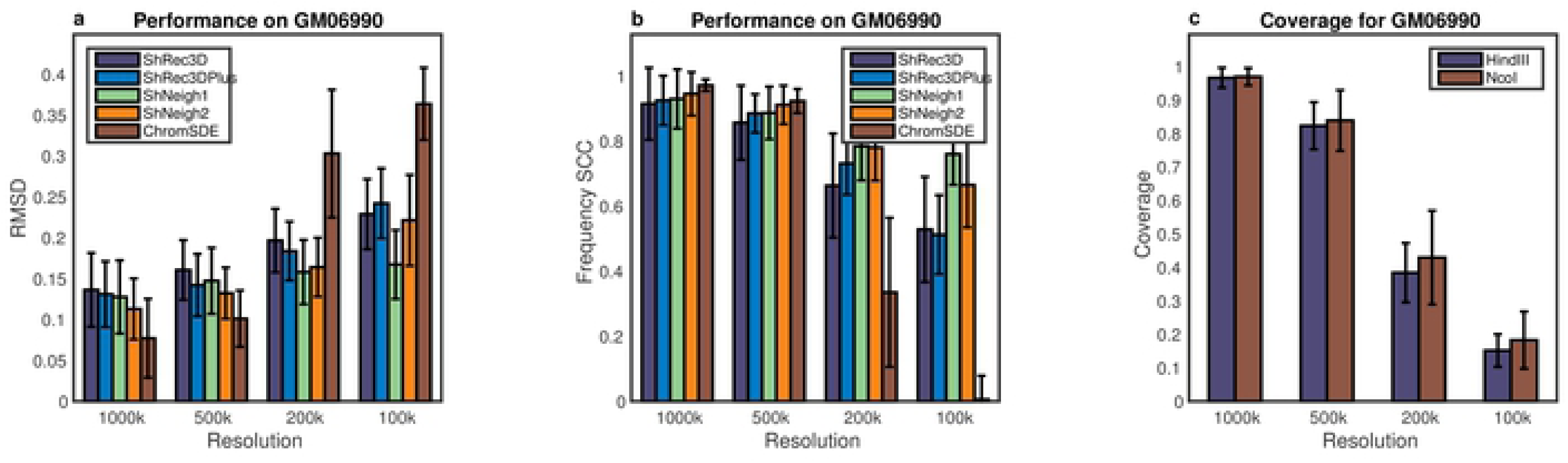
Performance comparison on GM06990 Hi-C data. (a) Spearman coefficient between the predicted distance matrices of HindIII enzyme and NcoI enzyme at different resolutions. (b) RMSD between the predicted structures of HindIII enzyme and NcoI enzyme at different resolutions. (c) Signal coverage of HindIII enzyme and NcoI enzyme at different resolutions. All measures are averaged across 23 chromosomes.

### Real Hi-C data study

As for the human GM06990 cell lines, we compute the average RMSD across 23 chromosomes (1-22 and X) between the predicted structures from HindIII and NcoI Hi-C data and the average Spearman correlation coefficient between the estimated distance matrices (dSCC) of the predicted structures from the two enzyme data, which are shown in Figure 5 and Supplementary Figure S3. Not surprisingly, all tested programs perform worse as the resolution rises (Figure 5a-b), since the average signal coverage gets lower at higher resolution (Figure 5c). We first compare the performances of ShRec3D and ChromSDE. Note that the average signal coverage is about 0.96, 0.82, 0.40, 0.17 for 1000k, 500k, 200k, 100k, respectively. The comparison between ShRec3D and ChromSDE shown in Figure 5 is very similar to the result shown in Figure 2 of Ref.[11]. We found the improvement of our shNeigh programs against ShRec3D is not so distinct as the case of simulated data at 1000k and 500k resolution, though the difference between them is still remarkable at 200k and 100k resolution. ChromSDE behaves the best at 1000k and 500k resolution but the worst at 200k and 100k resolution. The dSCC value of ChromSDE is even close to zero at 100k resolution (Figure 5b), reflecting that ChromSDE completely failed to recover the underlying structure of GM06990 data for very high resolution. On the contrary, ShNeigh1 and ShNeigh2 perform relatively stable across all resolutions, and shNeigh1 performs the best among all tested programs at 200k and 100k resolution. On the whole, the advantage of shNeig1 and shNeigh2 approaches maximum at high resolution but is limited at low resolution. Comparing Figure 5 with Figure 4, the advantage of ChromSDE shown at 1000k and 500k resolution seems that the noise level of GM06990 data is very low. However, we are more convinced by the conjecture that real Hi-C data are commonly the product of a mixture of diverse structures [2,24]. What’s more, the estimated conversion factor α by ShNeigh2 gets larger with increasing resolutions (Supplementary Figure S2), which coincides with the conclusion of Ref.[2].

The 3D structures of chromosome X predicted by ShRec3D, shNeigh1 and ChromSDE at different resolutions are drawn in Figure 6. At 1000k and 500k resolution, all three programs can give structures of relatively good reproducibility. However, at 200k and 100k resolution, only shNeigh1 generated clear and highly reproducible structures, while Shrec3D and ChromSDE just reconstructed some tangled messes.

**Fig 6.**
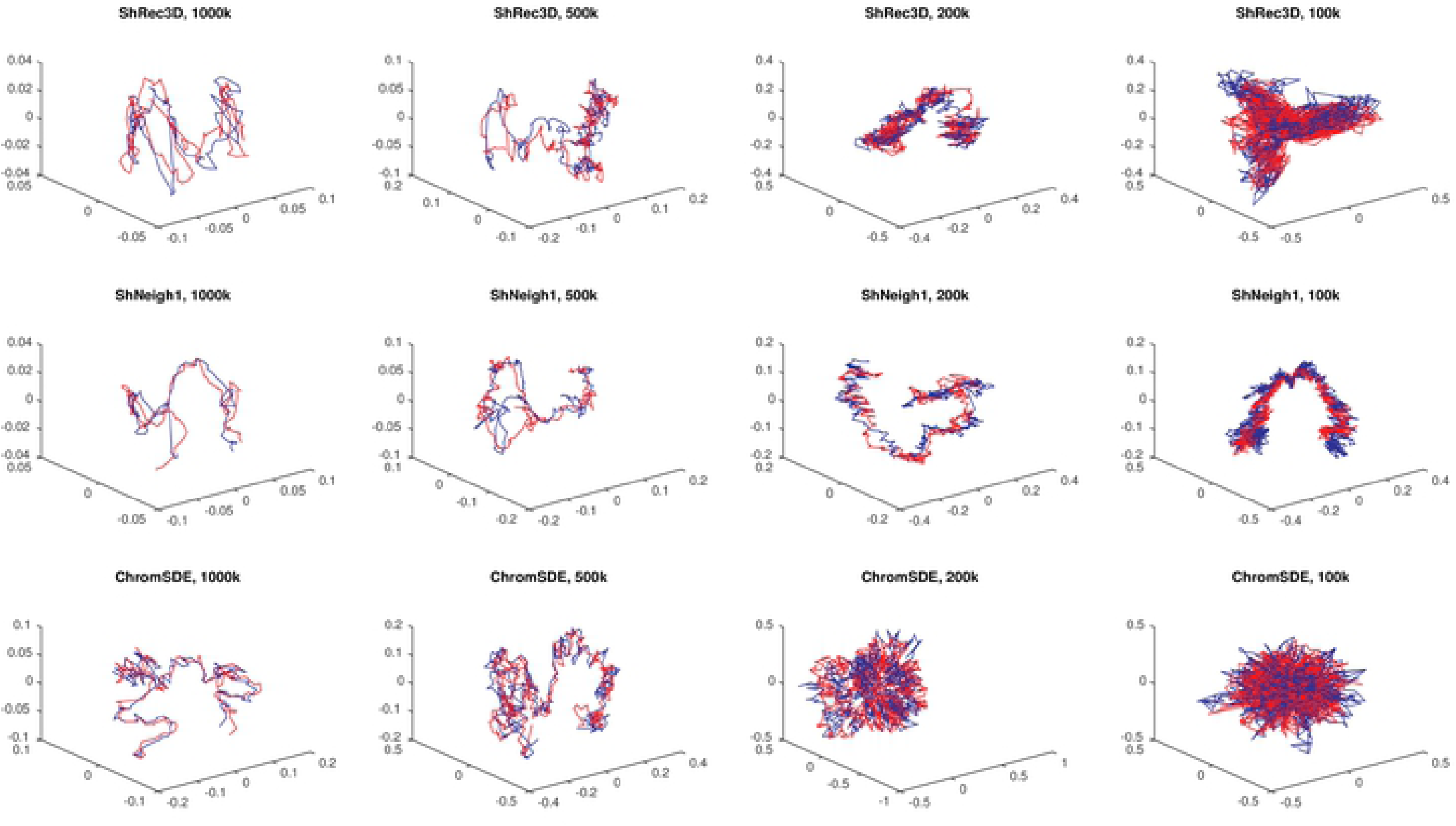
The alignment between predicted structures of chromosome X of GM06990 using HindIII enzyme(red) and NcoI enzyme(green) by ShRec3D, shNeigh1 and ChromSDE.

## Discussion

We have developed a novel method, named shNeigh, to reconstruct the 3D organization of chromosomes at the genome scale. It uses the classical MDS to minimize the gap between the predicted pairwise distances and those converted from the contact data. Shortest path algorithm is used to complete the converted distance matrix before applying MDS. ShNeigh explicitly models the local dependence of neighboring loci by a Gaussian expression and elaborately integrates the model into the MDS framework. Two strategies are adopted to determine the parameters (i.e. conversion factor α and the weight ρ) involved in the procedure: ShNeigh1 directly gives α=1 and ρ values by relating ρ with the loci number and the signal coverage, and ShNeigh2 searches for the two parameters through an iterative algorithm by minimizing the difference between the measured and predicted contact matrix.

Though ShNeigh2 has a step of searching for the optimal conversion factor α and the weight ρ, it is still much faster than ChromSDE. What’s more, our method achieves essential performance improvement compared to ShRec3D at some cost of time consuming (i.e. ShNeigh2) or even no time cost (i.e. ShNeigh1). Such an improvement exists and is quite apparent in most situations. Only for the data of high signal coverage that are generated from diverse structures the improvement gets somewhat weaker. Compared to ChromSDE, our programs are very robust in that they perform excellent for noisy or low signal coverage data, while ChromSDE works well for the data of low noise level and high signal coverage and corresponding to diverse structures. Considering it is very common for real Hi-C data to be noisy and sparse, our method is highly attractive. Observing Figure 5 and supplementary Figure S1, we conclude that ShNeigh can guarantee to obtain substantial improvement against both ShRec3D and ChromSDE for the Hi-C data with signal coverage not more than 0.5. On the contrary, the Markov chain that was used in HSA to model the local dependence of neighboring loci showed significant improvement only for very sparse Hi-C contact matrix (say, 10% signal coverage).

In this study, the proposed method combines the adjacency relationship of neighboring loci characterized by a Gaussian model with the classical MDS technique, and we have demonstrated this method can enhance the reconstruction quality of chromatin 3D structures significantly. We notice that our Gaussian adjacency model can easily be involved into most existing methods, including both distance based and likelihood based programs, such as HSA, PASTIS, ChromSDE, and so on. Assessing the performance of these various combinations is an interesting topic that deserves to be further explored in the future.

## Methods

One Hi-C experiment generates a library of paired-end reads. Each paired-end read represents one observation that the corresponding two restriction fragments contact each other. Then the reads are mapped to the reference genome and those of low quality are filtered out. After grouping the mapped high-quality reads according to genome loci where they locate, we get a contact frequency matrix *F*, where *F*_*ij*_ is a nonnegative integer representing the contact count between loci *i* and *j*. Here each locus is a genomic bin with a constant size such as 1Mbp or 40kbp. The resolution, namely the size of each genomic bin, is governed by the sequencing depth. The frequency matrix F is square and symmetric. Note that *F* may contain many zero entries generally, which indicates that the underlying locus pair are too far in the 3D space to interact with each other.

Given a frequency matrix *F*, our task is to reconstruct the 3D structure of the chromosome from which *F* is generated. That is, a coordinate matrix *X* = (*x*_1,_…,*x*_*N*_) ′ ∈ *R*^*N* × 3^should be derived from *F*, where *N* denotes the number of loci in the chromosome and *x*_*i*_ ∈ *R*^3 × 1^represents the 3D coordinate of the *i*-th locus. Our approach is based on the classical MDS methods.

### Classical MDS-based methods

Classical MDS-based methods, such as ShRec3d and ShRec3D+, generally consist of the following three steps.

First, convert the contact frequency matrix *F* into a distance matrix *D*. All existing methods, including MDS-based and likelihood-based approaches, assume that the contract frequency between two loci and their 3D distance agrees with the following power law relationship [2].

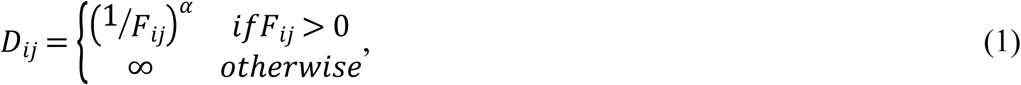

where α is the conversion factor and *D* _*ij*_ and *F*_*ij*_ are the 3D distance and contact frequency between loci *i* and *j*, respectively. The infinite distances *D*_*ij*_ = ∞ denote they provide no information for structure reconstruction. Eq.(1) does not consider the scale between the converted distance and the real physical distance. This scale, if necessary, can be described by adding a coefficient β before the term (1/*F*_*ij*_) ^*α*^ in Eq.(1). The parameter β is usually expressed explicitly in the objective function of likelihood-based methods. Our goal is to make the predicted structure align the underlying true structure as accurate as possible after applying scaling, reflection, translation and rotation operations, for which it is not requisite for β to emerge in Eq.(1). Thus, for the MDS-based methods β is calculated solely in assessing algorithm performance, namely in computing the RMSD criterion. The conversion factor α was set to a constant one in ShRec3D. Here we calculate α by the policy used in ChromSDE and ShRec3D+. See section 2.3 for detailed description.

Second, complete the distance matrix *D*. The classical MDS requires a full set of distances between all loci pairs available, but the infinite elements of *D* represent unknown distances and so must be endowed with finite values before applying MDS. To this end, we model the distance matrix *D* by a weighted graph whose nodes represent the genomic loci. In this graph two nodes i and j are linked by an edge if and only if the corresponding *D*_*ij*_ has finite value, and the length (or weight) of the edge is just the value of *D*_*ij*_. We define the distance between two nodes by the length of the shortest path relating them. Finding the shortest path between any two nodes in the graph is a classical problem in graph theory. As in ShRec3D, we use the Floyd-Warshall algorithm (implemented by the Matlab function *graphallshortestpaths)* to calculate the shortest paths and their lengths. Floyd-Warshall is a dynamic programming algorithm with time complexity *O*(*N*^3^), where *N* is the number of nodes. The resulting graph becomes a clique, namely a fully connected graph, and the shortest-path distances satisfy the triangular inequality. Note that some original finite distances may change their values after Floyd-Warshall calculation, reflecting the input data are noisy.

Third, map the distance matrix into 3D structure by multidimensional scaling. Multidimensional scaling (MDS) is a technique of data statistics that can determine the coordinates of *n* objects in the *k*-dimensional Euclidean space (here *k*=3) from their distance measures [22]. In order to elucidate the procedure of MDS, we firstly let *I*_*N*_denote an *N* × *N* unity matrix and **1** = (1,…,1)′ be a column vector of length *N* with all elements being ones, then we define an *N* × *N* matrix 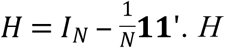 is symmetric and idempotent. Given the distance matrix *D*, construct the matrix 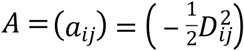 and further define *B* = (*b*_*ij*_) = *HAH*. Meanwhile, for the coordinate matrix *X* = (*x*_1,_…,*x*_*N*_)′ ∈ *R*^*N* × 3^ to be reconstructed we define its centralized inner product matrix by 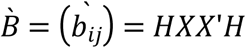. The classical MDS aims to minimize the following cost function:

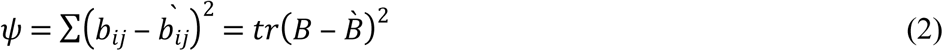

where tr(.) denotes the trace of a matrix. To this end, singular value decomposition is applied to *B* to get its three largest eigenvalues *λ*_1_*⩾λ*_2_*⩾λ*_3_ and their corresponding eigenvectors (*γ*_1_,*γ*_2_,*γ*_3_), with *γ*_*i*_ having been normalized to 1. Then the coordinate matrix *X* is recovered by

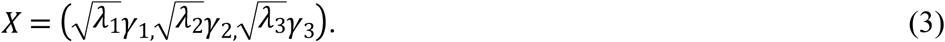

With this solution the cost function get minimum:

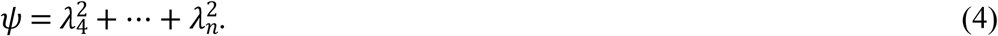

Therefore, when all eigenvalues other than the top three are equal to zero the reconstruction is exact. But in practice some *λ*_*i*_(*i*>3) may be negative, so the classical MDS can only approximately recover the chromosome structure generally.

### MDS with consideration of neighboring relationship

Intuitively, if two loci *x*_*i*_ and *x*_*j*_ are neighbors in the genome, the distance between the spatial coordinates of *x*_*i*_ and *x*_*j*_ should be small. In order to consider the local dependence of neighboring genomic loci, we define an affinity matrix *M* = (*m*_*ij*_) with

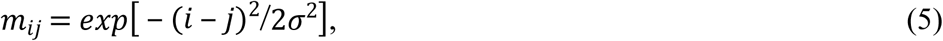

where s represents the rate that *m*_*ij*_decays with the genomic distance between loci *I* and *j*. Then we add the term cost to ∑*m*_*ij*_‖*x*_*i*_−*x*_*j*_‖^2^ into the cost function Eq.(2), turning the cost to

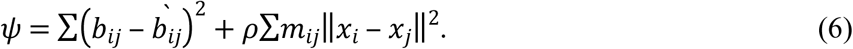

The second term reflects a distance penalty. It controls the smoothness of the reconstructed structure with a tuning parameter ρ. The extreme scenario ρ= 0 is just the ShRec3D [19] method, which gives a reconstruction entirely relying on the contact maps without smoothing.

After some algebra (see Supplementary text for a detailed derivation), we proved that the above problem is equivalent to minimizing the following object function:

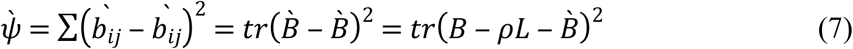

where 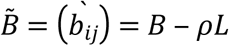, and *L* is the Laplacian matrix defined by *L* = *D* − *M* where *D* is the diagonal matrix with entries *d*_*ii*_ = ∑_*j*_*m*_*ij*_. Therefore, compared with Eq.(2), it is straightforward that we should exert singular value decomposition on 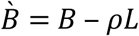 and get the top three eigenvalues 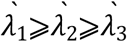 and their corresponding eigenvectors 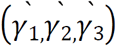 Then the reconstructed coordinate matrix *X* becomes

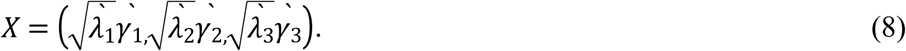

In the present work we only consider an affinity matrix M with the form of Eq.(5). Other forms of *M* are also desirable to attempt, for example, *m*_*ij*_ = 1 for |*i* − *j*| = 1 and *m*_*ij*_ = 0 otherwise. This matrix is just the scheme used in HSA [11], which captures the local dependency of the most neighboring loci solely.

### Parameter estimation

There are three parameters to be estimated in our method: the conversion factor α in Eq.(1), the distance penalty weight ρ in Eq.(6), the decaying rate σ in Eq.(5). Once their values are given, the reconstruction can be implemented by Eq.(8) straightly. We can either provide their values directly or infer them by an additional optimization procedure. We refer to the former as ShNeigh1 and to the latter as ShNeigh2.

For ShNeigh1, we empirically set 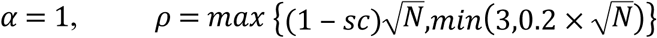 and *σ* = 0.023 × *N*, where *N* is the number of genomic loci and *sc* ∈ [0,1] denotes the signal coverage defined by the percent of non-zero entries in the contact matrix. *sc* is an indicator of the sparseness of the contact matrix. *α* = 1 is the policy adopted by ShRec3D. The expression of *ρ* is partly inspired by HSA. It means that the value of *ρ* is proportional to both one minus signal coverage and the root square of loci quantity. The term *min* 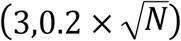 is used to handle the case of very high (close to 1) signal coverage. Without this term, *ρ* will tend to be zero as *sc* approaches 1. For ShNeigh2, we also set *σ* = 0.023 × *N*, but we infer α and ρ by minimizing an error function that describes the difference between the predicted frequency matrix 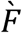 and the input frequency matrix *F.* Figure 1 gives a detailed description of the function *error* (*α,ρ,F*).

Minimizing *error*(*α,ρ,F*) with respect to α and ρ is a two-dimensional optimization problem, and it is difficult to calculate the gradient for *error*(*α,ρ,F*). ChromSDE used the golden section algorithm to optimize α, but it is a one-dimensional derivative-free algorithm and thus unsuitable for our context. Here we adopt the Nelder-Mead simplex (implemented by the Matlab function *fminsearch*), a multi-dimensional derivative-free algorithm, to simultaneously optimize α and ρ. A simplex in two dimensions is a triangular. For a given simplex, the Nelder-Mead simplex method first evaluates the objective function on its three vertices and recognizes the vertex with the largest value and the one with the smallest value. Then a new point with value lower than the vertex with the largest value is generated by operations of reflection, expansion and compression. A new simplex is thus constructed by substituting the largest vertex with the new point, or by shrinking toward the smallest vertex. Therefore, the minimum of the objective function can be approached by iteratively updating the simplex.

### Data

Both simulated and real Hi-C datasets are used to test the performance of our method. We generate the simulated datasets based on a helix curve structure with the following formula [2]:

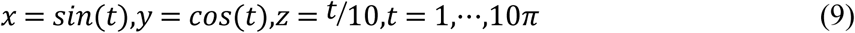

This structure is modeled by a linear polymer consisting of *N* points. The coordinates of the *N* points are calculated by Eq.(9) and then transformed to an *N* × *N* distance matrix *D*. In order to imitate the incompleteness nature of real Hi-C frequency matrix, only distances for *K* (*K*<*N*) nearest neighbors around each of the points are retained, and other distances are assigned to infinity. *K* directly determines the signal coverage of the transformed distance matrix *D* (see Fig.4f). The distance matrix *D* is then converted into the contact frequency matrix by 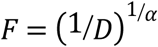 We further make the frequency matrix noisy by adding a random noise δ that is uniformly distributed in the region [-S,S], with *S* ∈ (0,1) being a given noise level. Specifically, 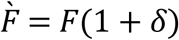. Finally the frequency matrix is scaled to summation 10^6^, which is similar to the usual treatment of real Hi-C data. Thus, the simulation code has 4 input parameters to be given by users: point number *N*, noise leverl *S*, conversion factor α and the number of nearest neighbors *K*. We fix the conversion factor *α* = 1throughout the simulation and tune the other three parameters according to different tasks.

There have been lots of in situ Hi-C data online, of which the human GM06990 cell dataset [1] is commonly used in literature. The advantage of this dataset is that it was generated with two different enzymes (HindIII, NcoI), making it possible to validate the structure of the investigated genome or validate alternative experimental designs. This dataset is also used in our present work. As described in the Introduction, the real Hi-C data need to be normalized to remove biases before reconstruction for all distance-based methods. The normalized contact frequency matrices of human GM06990 cells can be downloaded directly from the website of Amos Tanay’s group (http://compgenomics.weizmann.ac.il/tanay/?page_id=283). We compared our ShNeigh with three published programs: ShRec3D [19], ShRec3D+ [20] and ChromSDE [2], which are all distance-based methods, by using both the simulated and the real Hi-C data.

### Performance assessment measures

We use different structure similarity measures for simulated data and real Hi-C data to assess the performance of ShNeigh. Since the true structure is known for the simulated data, a natural measure is the Root Mean Squared Deviation (RMSD). RMSD measures the similarity of two structures by computing the distance of coordinates of the paired points between them. Given a real structure’s *N*×3 3D coordinates *P* = (*p*_1_,…,*p*_*N*_)′, and a predicted structure *Q* = (*q*_1,_…,*q*_*N*_)′(*p*_*i*_ or *q*_*i*_ is a 3 × 1 vector of the *i*th locus’ coordinate, i = 1, …, *N*), RMSD is defined as

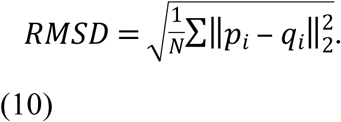

Before performing Eq.(10), some geometric operations: reflecting, rotating, translating and scaling, should be imposed on the predicted structure *Q* to make it align the true structure *P*. See [5,11, 23] for the detailed implementation. Obviously, smaller RMSD value means higher similarity of two structures and hence better performance of the tested program. It is widely used in bio-molecular structure comparison, such as protein structures and chromosome structures. In addition, we use the Spearman correlation coefficient (SCC) between the pairwise distances from the predicted structure and those from the true structure to give another performance measure.

As for the real Hi-C data, the underlying true structures of chromosomes are unknown, so the RMSD measure comes from comparing the two predicted structures of HindIII and NcoI enzymes. We also compute the Spearman correlation between the two estimated frequency matrices of the structures inferred from two different enzymes. It is more unbiased to use Spearman correlation than use Pearson correlation for testing every program, because Spearman correlation is independent of the conversion factor α [2]. Similar to Pearson correlation, the Spearman correlation value varies in [-1,1], the more close to 1.0 the better.

## Acknowledgments

This work was supported by the National Natural Science Foundation of China (No. 11604170, 61572286, 61873145), NSFC Joint with Zhejiang Integration of Informatization and Industrialization under Key Project (No. U1609218), Scientific Research in Universities of Shandong Province (No. J16LJ06) and the Natural Science Foundation of Shandong Province, China (No. ZR2019MA059, ZR2014AQ018).

## Supporting information

**S1 Fig.**
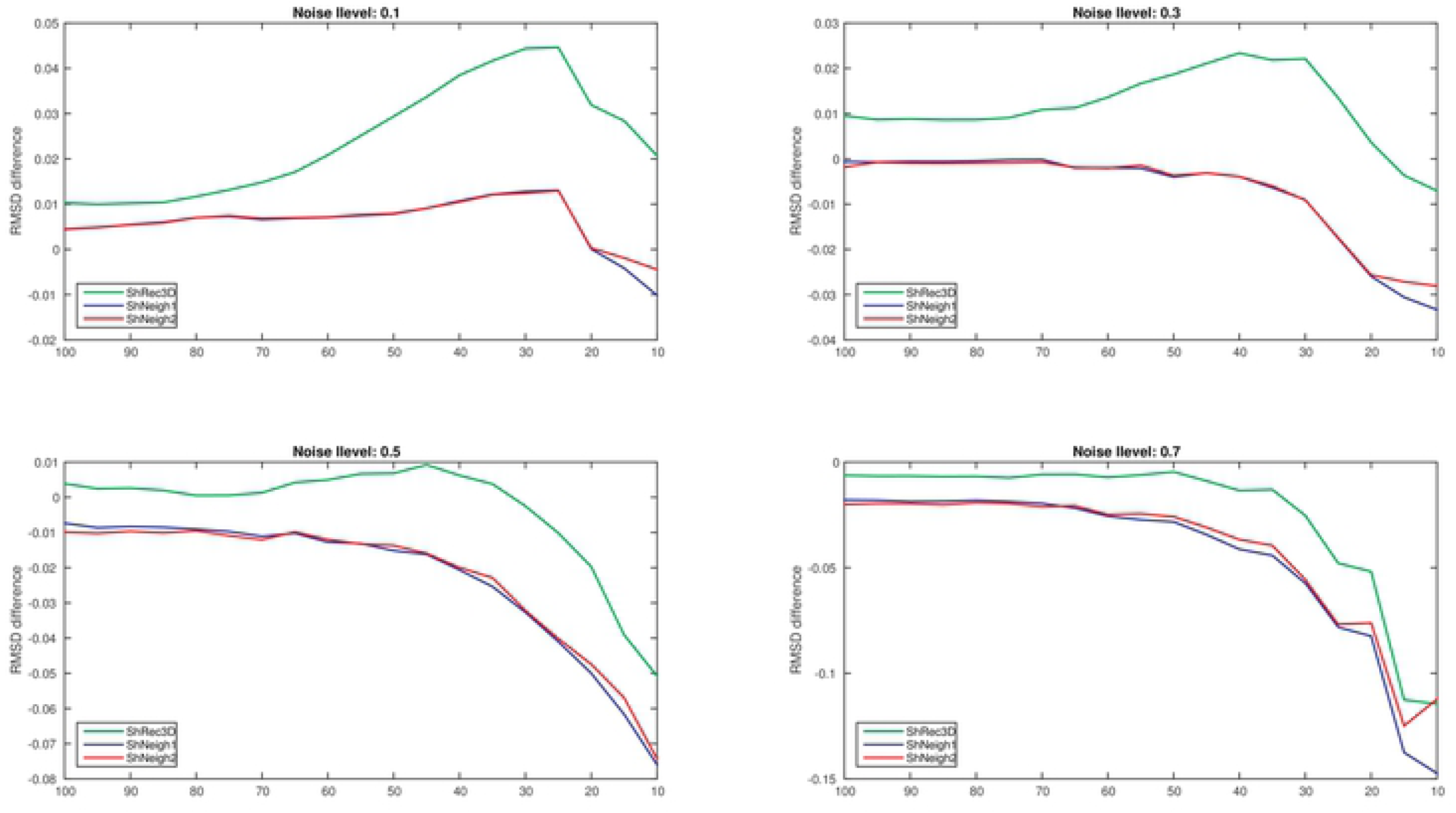
RMSD difference, i.e. RMSD of ShRec3D, ShNeigh1 and ShNeigh2 minus that of ChromSDE on simulated data under varying number of nearest neighbors K.

**S2 Fig.**
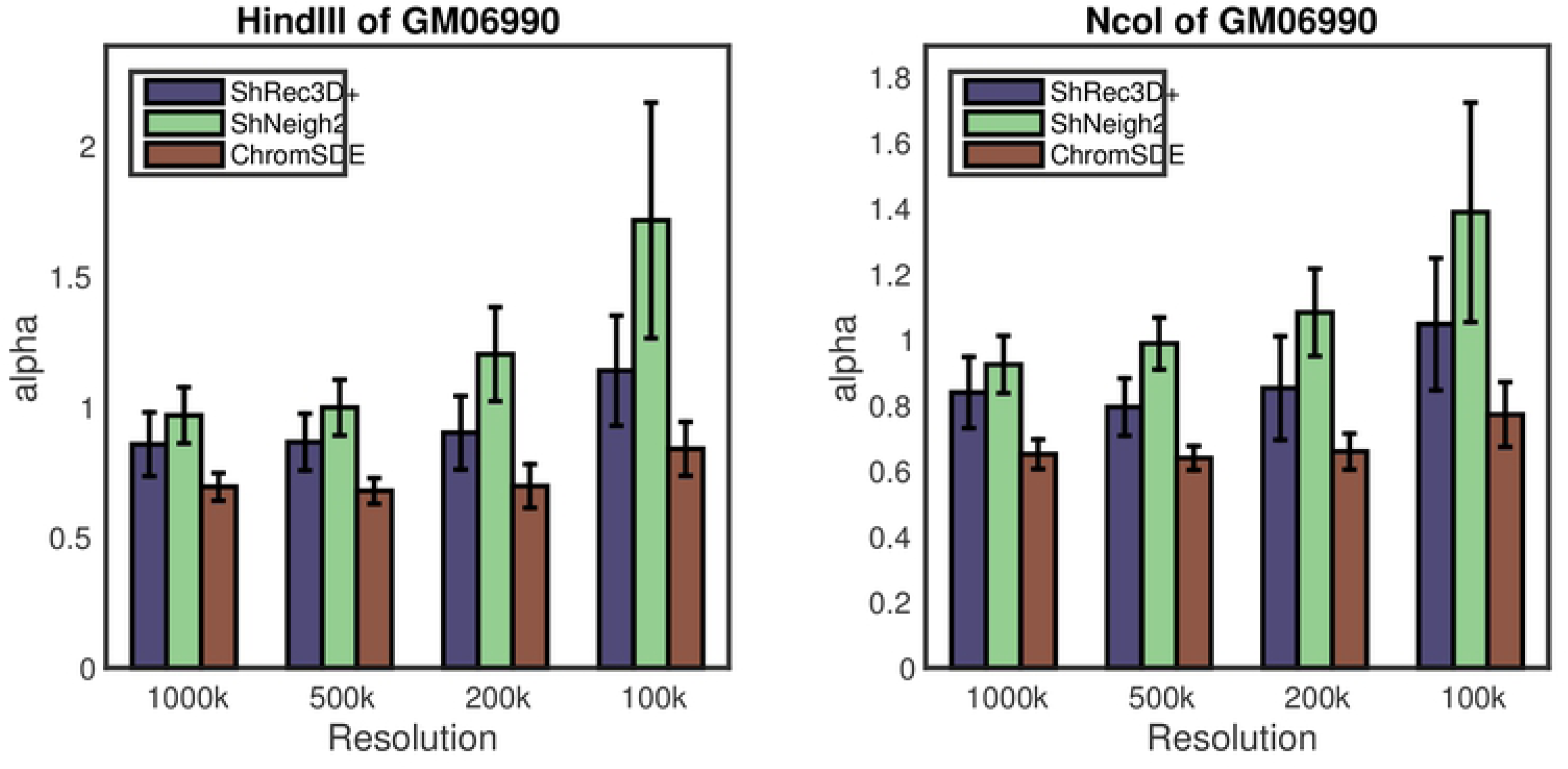
Estimated conversion factor of GM06990 Hi-C data at different resolutions for (a) HindIII enzyme and (b) NcoI enzyme. Averaged across 23 chromosomes.

**S3 Fig.**
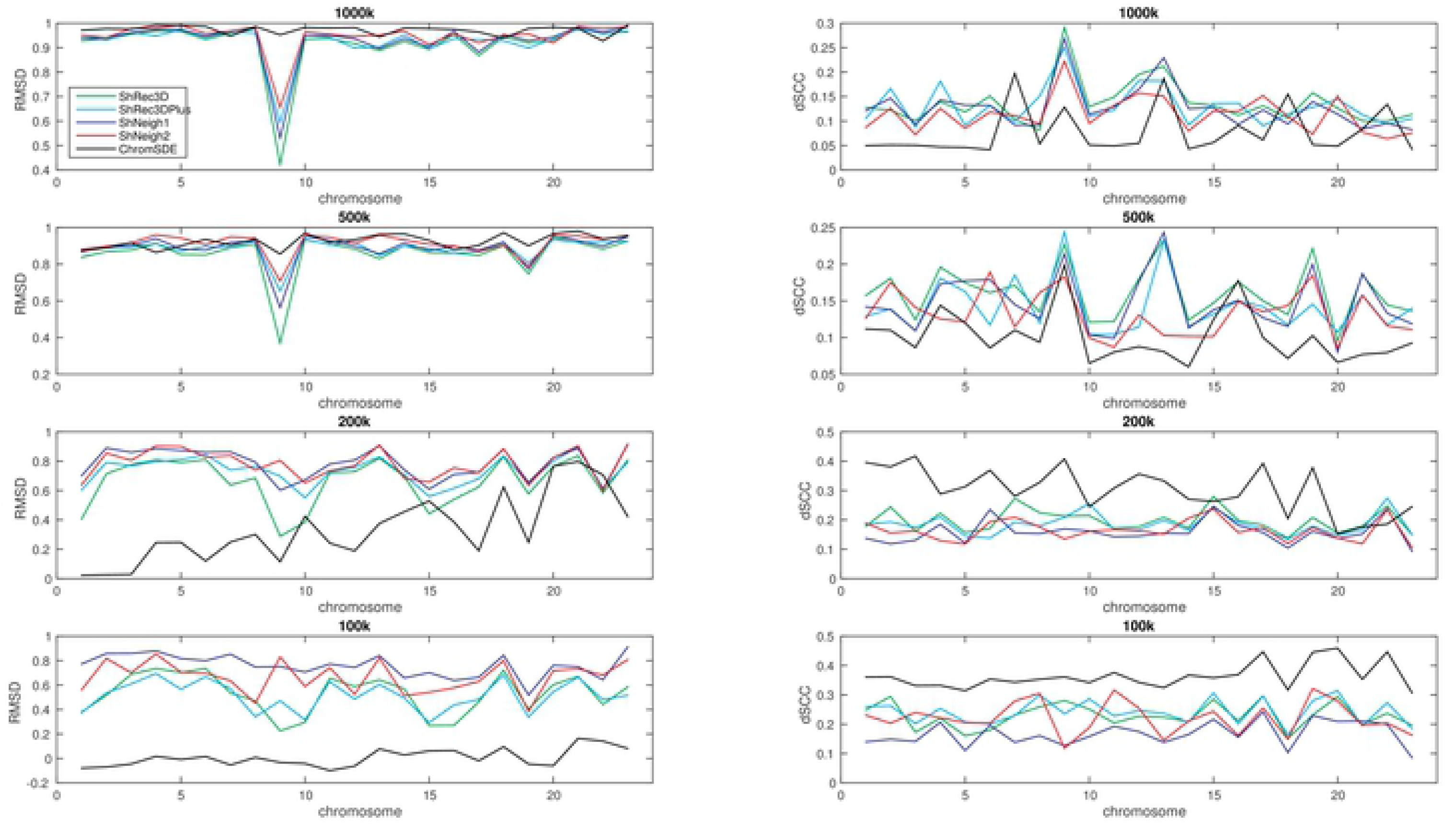
Performance of different methods on GM06990 Hi-C data at different resolutions for each chromosome. The left column corresponds to RMSD measure and the right column corresponds to dSCC measure, and each row represents one resolution.

**S1 File. supplement.docx**. Supplementary materials.

